# Detecting aquatic pathogens with field-compatible dried qPCR assays

**DOI:** 10.1101/2022.07.07.499119

**Authors:** Jessica Rieder, Pedro M. Martin-Sanchez, Omneya A. Osman, Irene Adrian-Kalchhauser, Alexander Eiler

## Abstract

Field-ready qPCR assays with a long shelf-life support monitoring programs for emerging aquatic pathogens and enable quick conservation and management decisions. Here, we develop, validate, and test the shelf-life of qPCR assays targeting *Gyrodactylus salaris* and *Aphanomyces astaci* with lyophilization and air-drying.

---

Pathogenic species are a major threat to aquatic and terrestrial ecosystems. Globalization (international trade, transportation, and urbanization) and anthropogenic global changes have been fostering the spread of pathogens (McIntyre et al., 2017; Guenard, 2021), resulting in biodiversity decline and economic losses. Three relevant aquatic pathogens with economic and ecologic implications are: (i) the monogenean salmon parasite *Gyrodactylus salaris* (*Gs*) that colonizes the skin, gills, and fins of salmon and has caused widespread losses in both wild and farmed Atlantic salmon (Bakke et al., 1992; Rusch et al. 2018), (ii) the crayfish pathogen oomycete *Aphanomyces astaci* (*Aa*) that elicits crayfish plague in native European, Asian, and Australian crayfish species and causes massive die-off events (Martín-Torrijos et al., 2021), and (iii) the amphibian-targeting panzootic chytrid fungus *Batrachochytrium dendrobatidis* (*Bd*), which originated in Asia, spread globally because of amphibian trade, and has decimated more than 500 amphibian species over the past half-century (Fisher and Garner, 2007, 2020; Scheele et al., 2019).

The analysis of environmental DNA (eDNA) is an emerging approach for quick, relatively inexpensive monitoring and detection of aquatic pathogenic organisms (Amarasiri et al., 2021). As a result, scientists, governmental agencies, and companies are increasingly incorporating eDNA methods into (semi)-automatic sampling machines coupled to portable real-time quantitative PCR (qPCR) thermocyclers for continuous on-site pathogen monitoring of waterways (Thomas et al., 2020; Sepulveda et al., 2019, 2020). However, a remaining challenge is the requirement of cold storage for key reagents, which prohibits their use in field-operating machinery. Reagents that can be dried and are stable at room temperature (RT) from are commercially available. However, they have not been independently evaluated for their applicability and true shelf-life regarding eDNA monitoring of pathogens.

This study describes field-ready storable dried qPCR assays for three aquatic pathogens, *Gs, Aa*, and *Bd*. The dried *Bd* assay was not evaluated for shelf-life, so results are not shown but worked upon reconstitution after drying. For *Gs* and *Aa* assays, we compared two different drying methods, lyophilization and air-drying, and amplification efficiency of dried assays across a time series (Table 1).

**Table 1.** qPCR assays evaluated in this study.

| Target<sup>a</sup> | Forward primers (conc.) | Reverse primers (conc.) | TaqMan probe (conc.) | IPC<sup>b</sup> | gBlocks fragments of reference sequences (Acc. No)<sup>c</sup> | qPCR program | Drying method | Shelf-life tested |
| --- | --- | --- | --- | --- | --- | --- | --- | --- |
| *Gs*<br>(Rusch et al., 2018) | Gsal-208F<br>(0.75 µM) | Gsal-149R<br>(0.75 µM) | Gsal-188P-MGB2<br>(0.25 µM) | Yes | Gs_124-289<br>(DQ898302) | 2 min 95°C;<br>45 cycles (10s 95°C, 1 min 60°C) | Lyophilization | Yes |
| *Aa*<br>(Vrålstad et al., 2009) | AphAstlT<br>S-39F<br>(1.2 µM) | AphAstl<br>TS-97R<br>(1.2 µM) | AphAstlTS-60T<br>(0.3 µM) | No | Aa_1-152<br>(AM947023) | 2 min 95°C;<br>45 cycles (5s 95°C, 20s 60°C) | Air drying | Yes |
| *Bd*<br>(Boyle et al., 2004) | ITS1-3<br>Chytr<br>(0.9 µM) | 5.8S-Chytr<br>(0.9 µM) | Chytr-MGB2<br>(0.25 µM) | Yes | Bd_26-271<br>(AY598034) | 2 min 95°C;<br>50 cycles (10 s 95°C, 1 min | Lyophilization | No |
|  |  |  |  |  |  | 60C) |  |  |

<sup>a</sup> Gs: *Gyrodactylus salaris*; Aa: *Aphanomyces astaci*; Bd: *Batrachochytrium dendrobatidis* <sup>b</sup> IPC: Internal Positive Control including a TaqMan probe plus its complementary standard DNA. <sup>c</sup>: gBlocks names refer to the selected positions in the corresponding reference sequences, whose GenBank Accession numbers are detailed

All three assays targeted the rDNA internal transcribed spacer 1 (ITS1) region and were evaluated for reproducibility and sensitivity in a wet, freshly-made state. The standard curves were generated using serial dilutions of synthetic double-strain DNA fragments (gBlocks, Integrated DNA Technologies, Inc., Belgium) encompassing the target regions of the three assays (Table 1; Fig.1a,c; Supp. Material Fig. S1).

**Figure 1.**
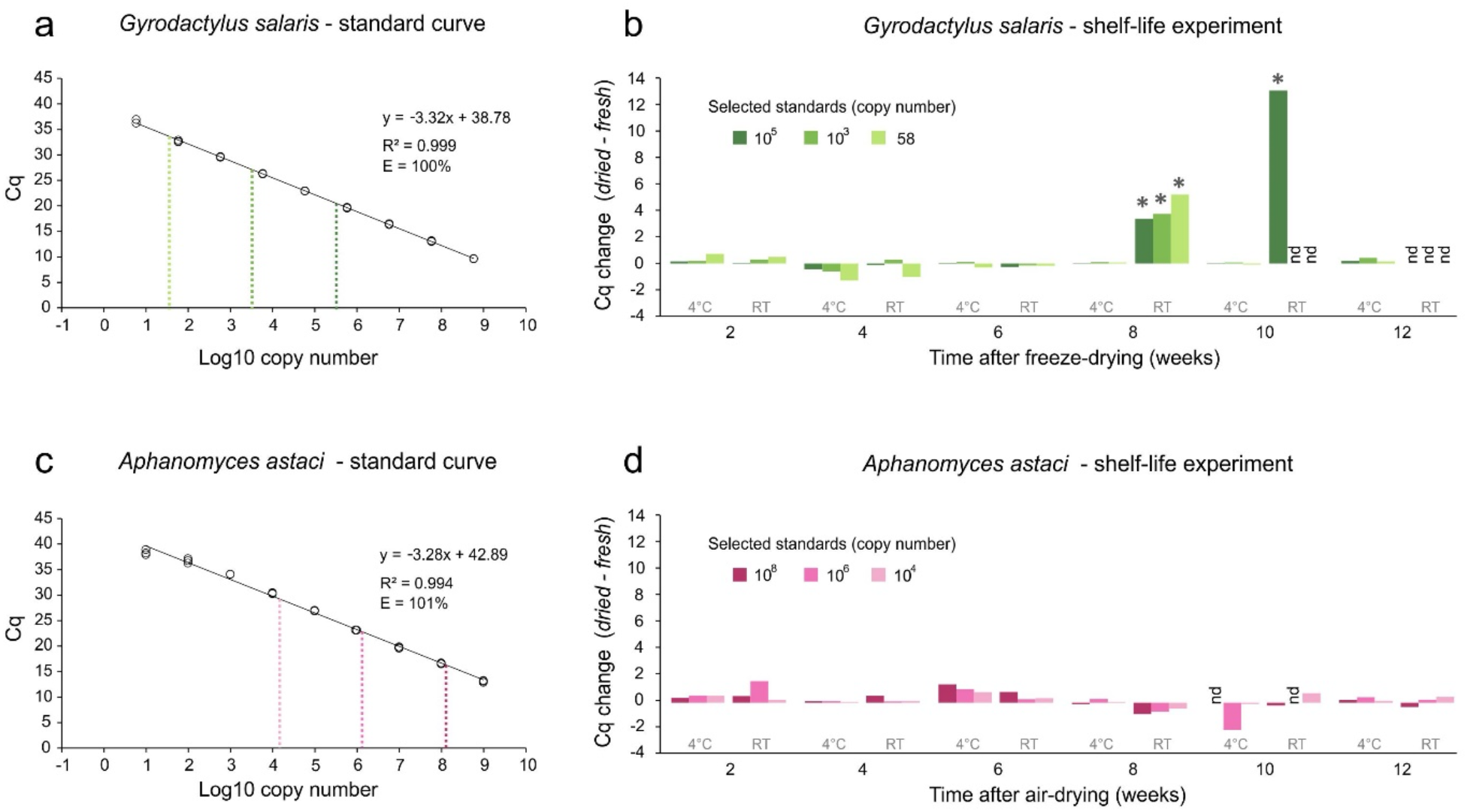
Validation and stability results for the two dried assays. (a + c) Standard curves of TaqMan-based qPCR amplification of *Gyrodactylus salaris* (*Gs;* a) and *Aphanomyces astaci* (*Aa*; c) using fresh assays and gBlocks fragments. Standard curves were plotted using all three replicates for each serial dilution. The dotted lines represent the three concentrations used in each shelf-life experiment. (b + d) Shelf-life experiment results for *Gs* (b) and *Aa* (d) over 12 weeks, testing three concentrations and two different storage temperatures (4°C and room temperature - RT). These results are shown as changes in Cq values compared to fresh assay (Cq dried – Cq fresh). Concentrations for each assay were selected within the quantification range. Asterisks indicate the main Cq changes associated with the degradation of the assays (see details in Fig. 2).

After generating baseline data for the wet assays, the efficiency and shelf-life of dried assays for *Gs* and *Aa* were evaluated with a 12-week time-series experiment. The *Gs* assays were prepared using SensiFAST Lyo-Ready (Meridian Biosciences, Bioline Assays Ltd, UK) with an exogenous internal positive control (IPC; Applied Biosystems), which allows for the assessment of both the overall integrity of assays and the potential false negatives (PCR inhibition) in future environmental analyses. *Gs* assays were frozen at −80°C for 24h and then lyophilized at −50°C and <0.1 mbar for 4h with a FreeZone 2.5 Liter Benchtop (Labconco, USA). *Aa* assays were prepared with Air-Dryable qPCR Mix (Meridian Biosciences, Bioline Assays Ltd, UK) and air-dried at 65°C for 65 min using a drying oven with a fan speed of 100%; no IPC was used (Table 1). Both assays were vacuum-sealed in bags with silica beads, placed in darkness, and stored at either 4°C or RT (21°C ±1°C). qPCR analyses comparing dried vs. fresh assays were conducted every two weeks post-drying. Three different concentrations of the gBlocks fragments were used as standards for *Gs* (5.8 × 10^5^, 5.8 × 10^3^ and 58 copies of Gs_124-289) and *Aa* (1.9 × 10^8^, 1.9 × 10^6^ and 1.9 × 10^4^ copies of Aa_1-152) (Fig. 1).

We find that in three of four conditions (i.e., *Gs*: 4°C, *Aa*: 4°C, RT), dried assays perform equally well as fresh assays even after 12 weeks (3 months) of storage. Only *Gs* assays stored at RT declined in performance at week 8, with increased Cq values and anomalous IPC signals (Fig. 1b, indicated by the asterisks; Fig. 2). In optimum conditions, the Cq values for IPC (VIC) should be 25±2, as shown in Fig. 2b for the assays stored at 4°C and fresh controls. While at week 10, only the highest concentration could be detected, by week 12, all concentrations were undetectable (Fig. 1b). Since the aim was to develop *Gs* assays stable at RT, further optimization is required to make this assay stable at RT for the same duration. In a diagnostic setting, however, often a combination of storage options is possible, where assays can be stored for a longer time at 4°C and used or stored at RT for field-based studies (<6 weeks) when cold storage is not possible. Air-dried *Aa* assays were stable until the end of the experiment at all concentrations and in both storage conditions. An anomaly occurred in week 10 when the highest concentration of the 4°C stored group and the medium concentration of the RT stored group were not detected. Since results in the following timepoint, in week 12, were on par with the control group, we assume that this anomaly was human or machine-based.

**Figure 2.**
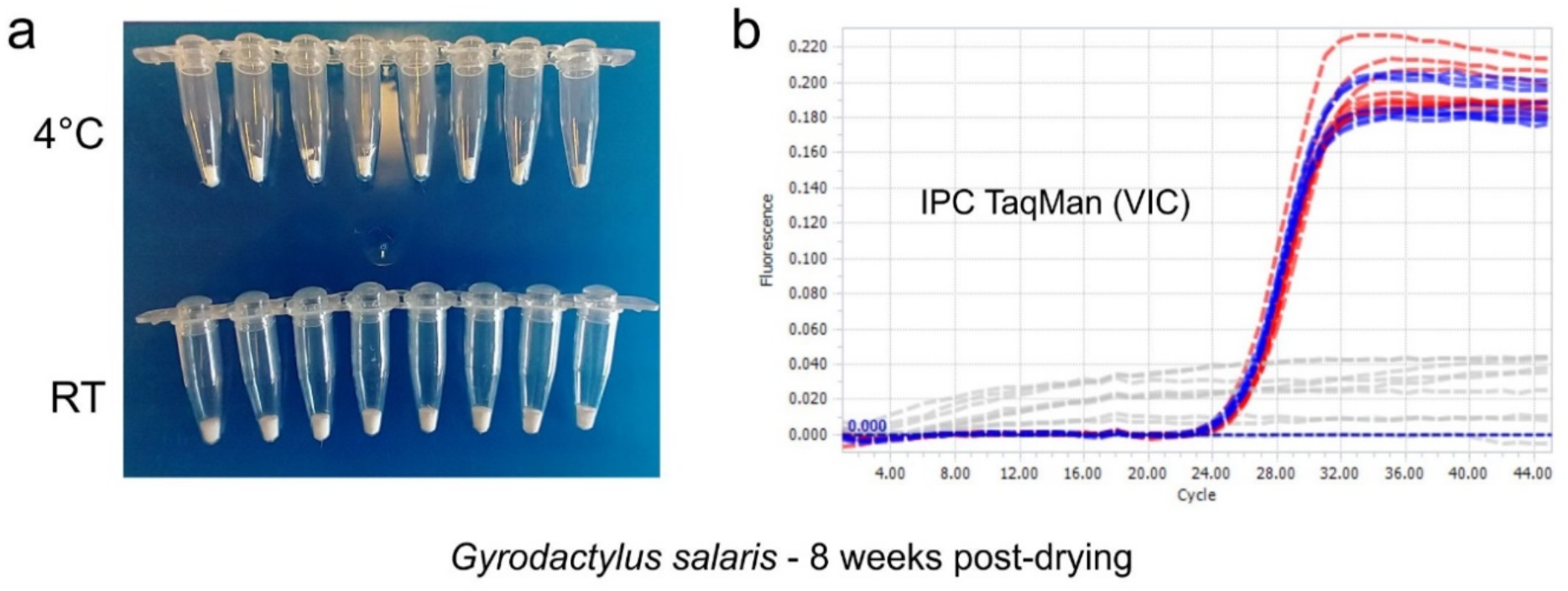
Partial degradation of the freeze-dried *Gyrodactylus salaris* qPCR assays. (a) Aspect of qPCR reagents. (b) Amplification curves for the internal positive controls (IPC; VIC signals). Note the poor performance of the dried assays stored at room temperature (grey IPC curves) compared to those stored at 4°C (red) and fresh controls (blue).

The development of field-ready diagnostic assays is vital for detecting and controlling emerging diseases quickly on site. Here, we provide proof-of-concept data for field-ready qPCR assays that could be further coupled with portable field-use qPCR machines to detect and monitor aquatic pathogens. Additional steps include further optimization to increase shelf-life and easy transferability to developing (semi)-automatic microfluidic devices. A possible method for ease of transferability would be to follow Xu et al. (2021), where the addition of liquid nitrogen to the master mix formed a transferable ball.

We demonstrate the feasibility of preparing dried, long-term stable qPCR reactions that can be reconstituted with water and a template. All assays would be suitable for short-term field-based conservation monitoring programs.

This work was supported by the Norwegian Environment Agency (Auto e-DNA project).

## Author statement

Rieder: conceptualization, methodology, validation, writing - original draft, and writing – review & editing Martin-Sanchez: conceptualization, methodology, investigation, writing - original draft, and writing – review & editing

Osman: methodology, writing – review & editing

Adrian-Kalchhauser: funding acquisition, supervision, writing – review & editing

Eiler: conceptualization, funding acquisition, supervision, writing – review & editing

