## Supplemental Material Figure S1 for "Detecting aquatic pathogens with field-compatible dried qPCR assays"

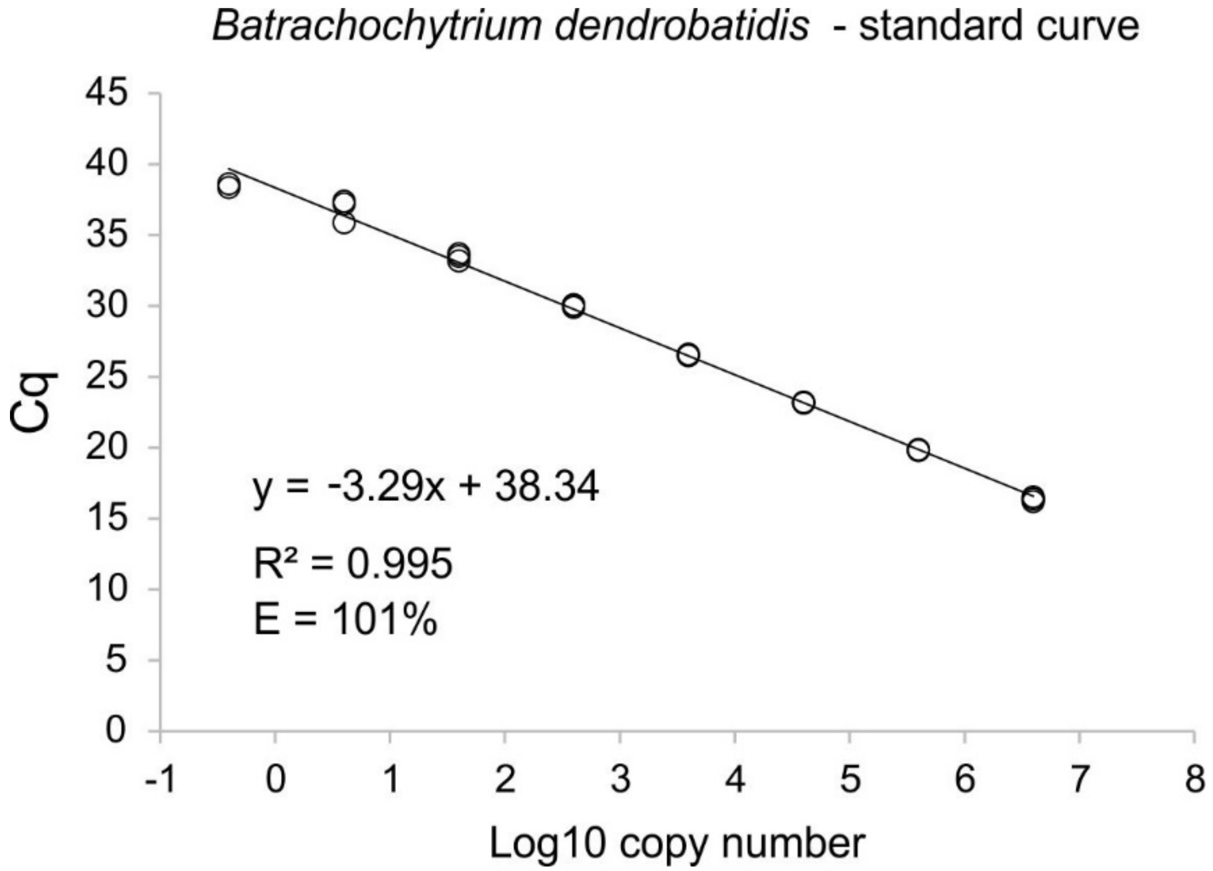


**Figure S1**. Standard curve for the *Batrachochytrium dendrobatidis* (Bd) TaqMan assay using fresh reagents. This includes all three replicates for each serial dilution of the gBlocks fragment Bd_26-271.
